# Quantitative Modeling of the Short-Term Response to Nitrogen Availability that Coordinates Early Events in Lateral Root Initiation

**DOI:** 10.1101/2023.12.05.570292

**Authors:** Allison Gaudinier, Lisa Van den Broeck, Miguel Moreno-Risueño, Joel Rodriguez-Medina, Rosangela Sozzani, Siobhan M. Brady

## Abstract

Nitrogen (N) is an essential macronutrient and its bioavailability plays a major role in how plant development is tuned to environmental nutrient status. To find novel factors in early root system architecture responses to N conditions, we performed *Arabidopsis thaliana* root transcriptome profiling of a short-term time course in limiting and sufficient N conditions. Using this data, we inferred transcriptional regulatory networks in each condition, which revealed the N-condition specific responses of jasmonate regulation; transcriptional factor (TF) ERF107 plays a more generalized role in lateral root development while TF LBD13 is specific to N-limiting conditions. Further, we used a single cell LR cell-type specific transcriptome dataset to model and analyze the roles of TFs LBD13, ERF107, and PDF2 in early stages of LR development. Linking the N time course transcriptomics, LR mutant phenotypes, and cell-type specific single cell profiling, these approaches provide multiple lines of evidence to find and test the roles of TFs that are involved in early root patterning responses to N conditions.

## Introduction

Nitrogen (N) is an essential macronutrient for plants. However, its bioavailable levels are often insufficient, limiting plant growth and productivity. The sensing and uptake of N plays a crucial role in plant physiology and contributes to the regulation of many developmental and metabolic processes. As such, N is added via fertilizer to ensure sufficient plant growth and yield, however, this results in extensive downstream negative consequences to the environment.^1^ In the model species *Arabidopsis thaliana*, changes in N availability have a rapid impact on the transcriptome; many genes involved in N uptake and assimilation are upregulated within minutes of exposure to higher N concentrations.^2–4^ These rapid responses allow for plants to take advantage of this potentially transient nutrient source. Additionally, N availability shapes root system architecture (RSA), largely by changes in lateral root initiation or elongation to scavenge N in a limiting environment.^5–7^ Changes in RSA are a common response to altered nutrient status in the environment, and allow the plant to tune its investment in root growth with its needs for nutrient acquisition.^5^ The factors that regulate these early transcriptional changes which lead to N-responsive developmental changes in RSA are largely unknown and those described do not sufficiently explain the N response.

Transcriptome datasets that profile expression in response to altered N conditions of ammonium and/or nitrate are diverse in their experimental design (different N concentrations and sources, and tissues sampled) and are collectively complementary as a data source to identifying genes that regulate N-mediated growth and metabolism. Temporal sampling of plants exposed to altered N conditions revealed rapid and dynamic expression changes for N transporters, metabolic genes, and TFs.^2–4^ These datasets, along with additional *in vitro* and *in vivo* data, can serve as input for network inference.^7–14^ Complex and interconnected transcriptional regulatory networks have been revealed^15^, including the identification of six transcription factors (TFs) as important regulators of N uptake and metabolism CIRCADIAN CLOCK ASSOCIATED 1 (CCA1), AUXIN RESPONSE FACTOR 8 (ARF8), BASIC LEUCINE-ZIPPER 1 (BZIP1), the TGACG SEQUENCE-SPECIFIC BINDING PROTEIN 1/4 (TGA1/4) double mutant, and CYTOKININ RESPONSE FACTOR 4 (CRF4). ^4,16–19^ Furthermore, *bzip1* and *tga1/4* have RSA defects due to a misregulation of N metabolism, linking N-mediated changes in transcriptional regulation to root development. There is insufficient data, however, connecting short-term N exposure and its link with development.. In this study, we addressed a knowledge gap through short-term transcriptional profiling of Arabidopsis roots in response to variations in N concentration. Network inference and modeling were employed to uncover additional transcriptional regulators of N-responses that contribute to the developmental regulation of RSA. Our findings demonstrate that changes in nitrogen availability have a significant impact on transcriptional networks, notably those linked with lateral root development and jasmonate (JA) signaling. *CCA1* was confirmed as a transcriptional regulator of N metabolism, while LOB DOMAIN-CONTAINING PROTEIN 13 (LBD13) emerged as a central regulator of N-mediated regulation, influencing lateral root emergence in a context-specific manner. The interaction of LBD13 within a feedforward loop with PROTODERMAL FACTOR 2 (PDF2) and ETHYLENE RESPONSE FACTOR 107 (ERF107) was quantitatively modeled to simulate these regulatory interactions during early lateral root initiation. The model highlighted the importance of *LBD13’*s spatiotemporal expression for proper transcriptional progression in this critical mode by which roots alter their development during changes in N availability.

## Results

### Dynamics of Nitrogen-Responsive Transcriptional Networks show Crosstalk with Jasmonate Signaling in Arabidopsis Roots

As N is added to augment its limited bioavailability in the soil via fertilizer, we sought to characterize how a plant’s transcriptome changes upon a sudden increase in N concentration. *Arabidopsis thaliana* seedlings (Col-0) were grown for seven days on media with limiting nitrogen (1mM KNO_3_) and then transferred to either limiting (1mM) or sufficient (10mM) KNO_3_. Roots were collected immediately (0 minutes used exclusively for the 1mM KNO_3_ condition as a control for mechanical transfer), 15, 45, 90, and 180 minutes after transfer (Figure 1A). To identify transcriptionally responsive genes to this change in KNO_3_ concentration, we performed pairwise comparisons at each of these time points between N sufficient and N limiting conditions (q < 0.05). A total of 736 genes were detected as differentially expressed (DEGs), the majority of which showed distinctive and dynamic expression patterns. Specifically, 698 DEGs (94.84%) showed differential expression at only one time point (Figure 1B) and 38 DEGs (5.16%) were shared across at least two time points (Supplemental Figure 1, Supplemental Table 1). Such diversity in temporal transcriptional responses in response to a sudden increase in N suggests complex underlying regulatory networks.

**Figure 1.**
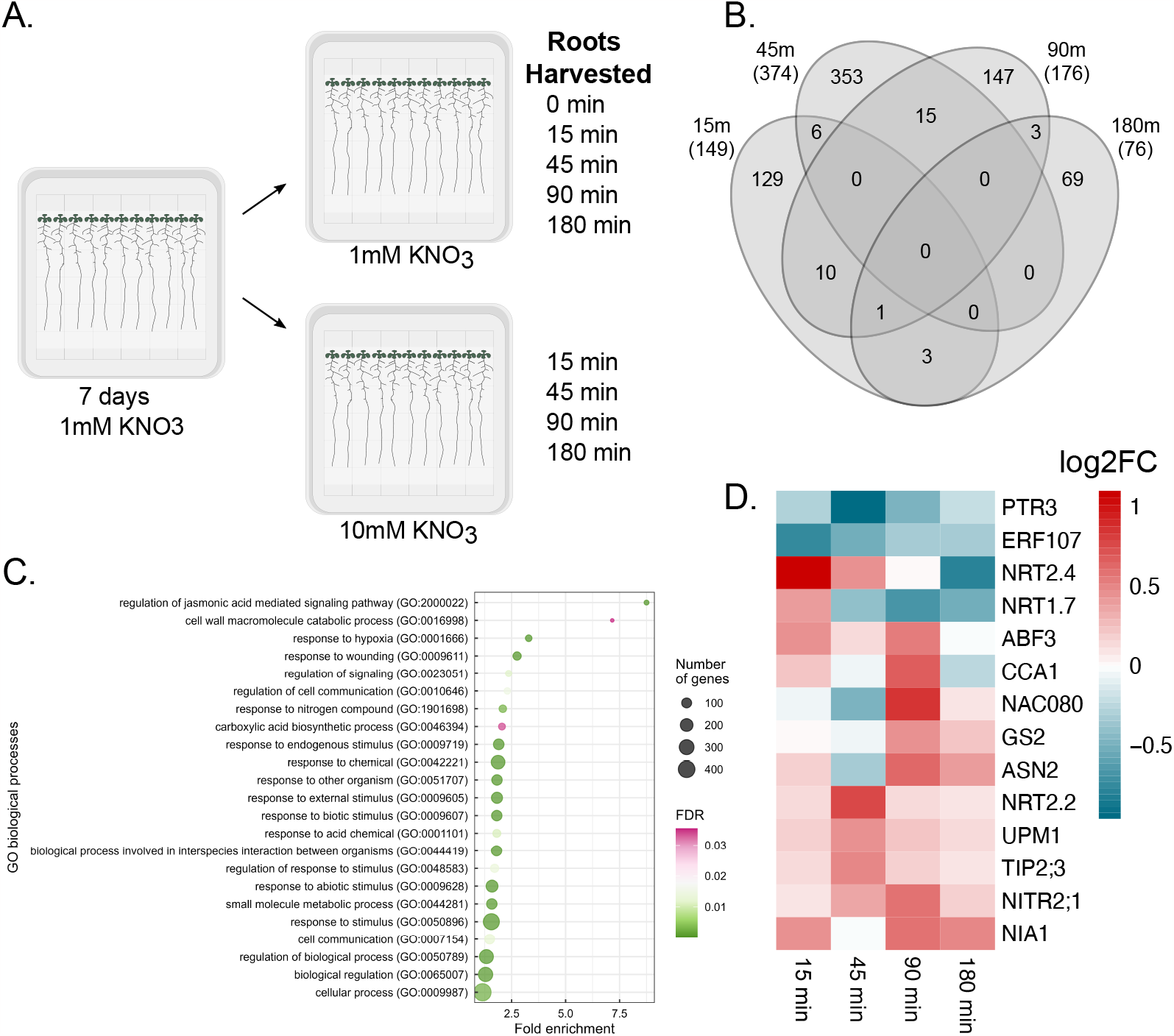
Transcriptional response upon nitrogen acclimation. **A**. Experimental design of the time course. **B**. Number of differentially expressed genes (DEGs) at each time point comparing nitrate sufficient with limiting conditions (q < 0.05). **C**. Gene Ontology enrichment of DEGs. **D**. Heatmap of differentially expressed genes linked to N.

To validate that our experimental design captured canonical aspects of the N response, we conducted enrichment analyses. When comparing our 736 DEGs against three diverse but complementary datasets that capture aspects of N-mediated transcriptional regulation, we found significant enrichment for N-responsive genes. Firstly, we found six DEGs in common with the fifty most broadly responsive N genes identified in a meta-analysis study^20^ (Fisher exact p-value = 0.00211). In a comparable expression profiling dataset^21^, we found 27/474 genes in common (Fisher exact p-value = 0.00017). Finally, we found an enrichment of our DEGs in our previously published yeast one-hybrid (Y1H) transcriptional network^7^ for N- and N-associated metabolism^6^ with 23/431 genes (Fisher exact p-value = 0.00126) (Supplemental Table 2A-C). Additionally, to further validate the robustness of our dataset, we performed a gene ontology (GO) enrichment and found that ‘response to nitrogen compound’ (GO:1901698) was significantly enriched within the DEGs (Figure 1C). When exploring the expression profiles for several nitrate transporters, nitrate assimilation genes, and known nitrate-related transcription factors, we found that eleven of these genes showed a time point-specific induction, while three DEGs were repressed, highlighting the diverse temporal transcriptional nature of N-dependent responses (Figure 1D).

We then hypothesized that distinct transcriptional cascades result in a unique molecular response triggered at each time point upon differential nitrate availability. To identify transcriptional regulators enabling such a hierarchical response, we inferred gene regulatory networks (GRNs) with the previously identified 736 DEGs (FDR < 0.05), of which seventy-three are transcription factors (TFs) (Supplemental Table 3A). We inferred two networks, one to explore regulatory interactions in nitrate limiting conditions, and the other in nitrate sufficiency. Changes in wiring (interactions between TFs and their downstream targets) were inferred using a machine learning approach^22^ (Supplemental Table 3B,C) (see Methods). The network in N-limiting conditions contained 277 genes and 382 interactions, while the N sufficient network comprised 279 genes and 384 interactions (Supplemental Figure 2A,B). In 28% of these inferred interactions, we found that the *cis*-elements present in the promoter of the target gene were bound by their respective TF *in vitro*, providing validation of our approaches^11^ (Supplemental Table 4). When comparing these two networks (Figure 2), major changes in connectivity were observed between JA signaling genes.

**Figure 2.**
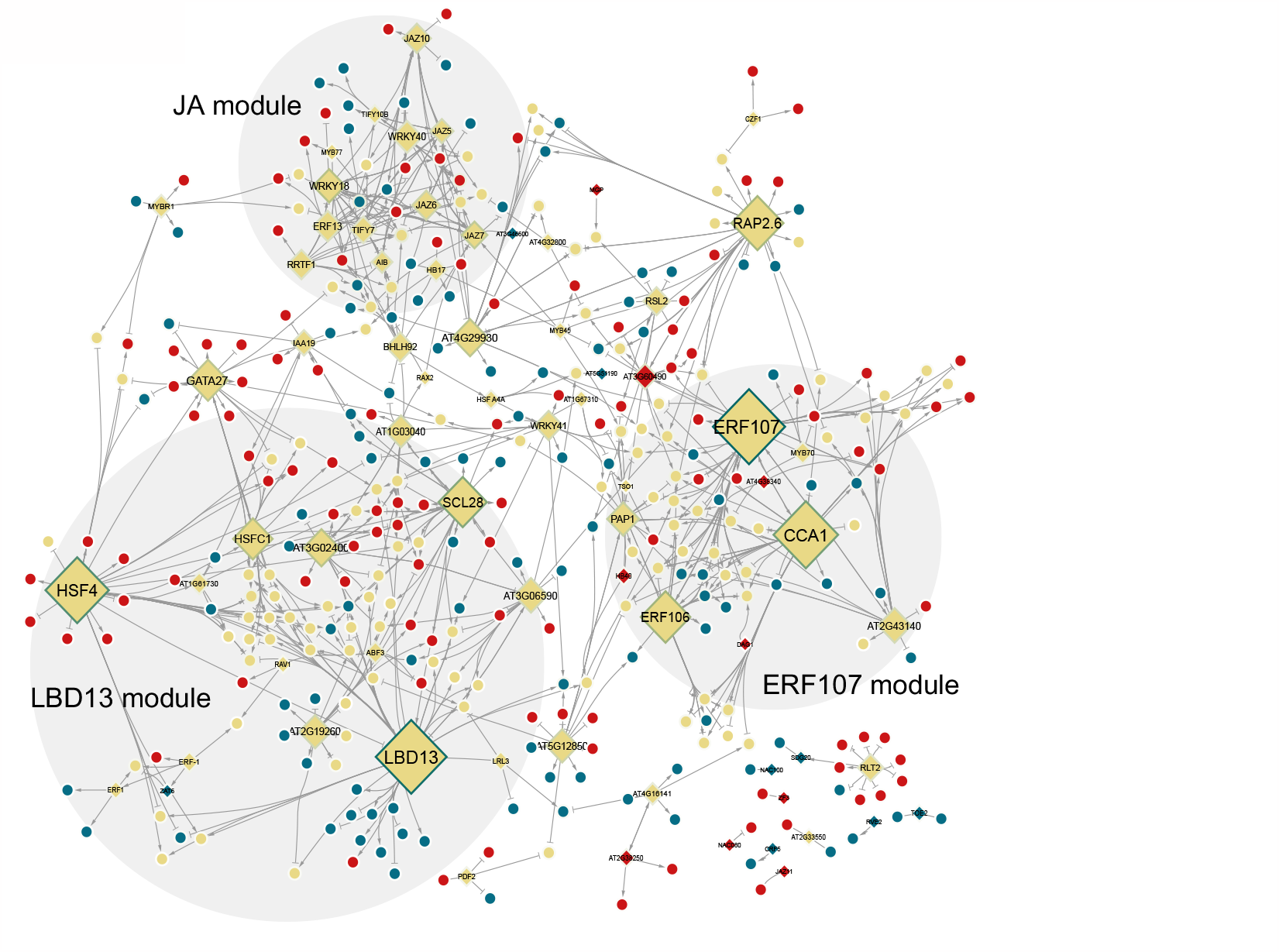
Network displaying the dynamical rewiring upon KNO_3_ exposure. The border color represents the DyNet rewiring score (darker is a higher score). Yellow nodes are genes in 1mM KNO_3_ and 10mM KNO_3_ networks, Blue nodes are in the 1mM network only, Red nodes are in the 10mM network only. Activating and repressing interactions are represented by point and block arrows respectively.

Within this set of 736 DEGs used to infer the two networks, JA signaling (‘regulation of jasmonic acid mediated signaling pathway’ [GO:2000022]) was the most significantly enriched GO category (Figure 1C, Supplementary Figure 3 [heatmap for all JA genes]); and 171 of these genes were previously identified to be transcriptionally responsive to JA^23^ (Fisher exact p-value = 4.974e-13, Supplementary Table 2D). These genes include the JA precursor biosynthesis genes LIPOXYGENASE 3 (LOX3)^24^ and OXOPHYTODIENOATE-REDUCTASE 3 (OPR3)^25^, the JA catabolic enzyme JASMONATE-INDUCED OXYGENASE4 (JOX4)^26^, as well as JA signaling repressors, including eight JASMONATE-ZIM-DOMAIN PROTEIN (JAZ) proteins^27^, and the TFs WRKY DNA-BINDING PROTEIN 18 (WRKY18) and WRKY40^28^. Most of these JA signaling DEGs have reduced expression in elevated N, specifically at the 15 minute time point (Figure 3C, Supplemental Table 1). In addition to this over-representation, JA-associated genes form a highly interconnected module in both the limiting and sufficient nitrate inferred networks (Figure 2A). Despite their annotation as JA-associated genes, only 11 of the total 99 interactions across these modules are in common between the two networks (Figure 3A-B). Interactions with likely functional consequences include predicted targets of the JAZ transcriptional repressors (JAZ2, 5, 6, 7, 9, 13), as well as WRKY18 and 40 which are vastly rewired in the two N networks. This suggests dynamic regulation of JA signaling in roots exposed to elevated N concentrations. To our knowledge, this is the first time that the JA-response has been linked to N responses through transcriptomics. JA pathway genes have been shown to affect lateral root growth. ^29–33^

**Figure 3.**
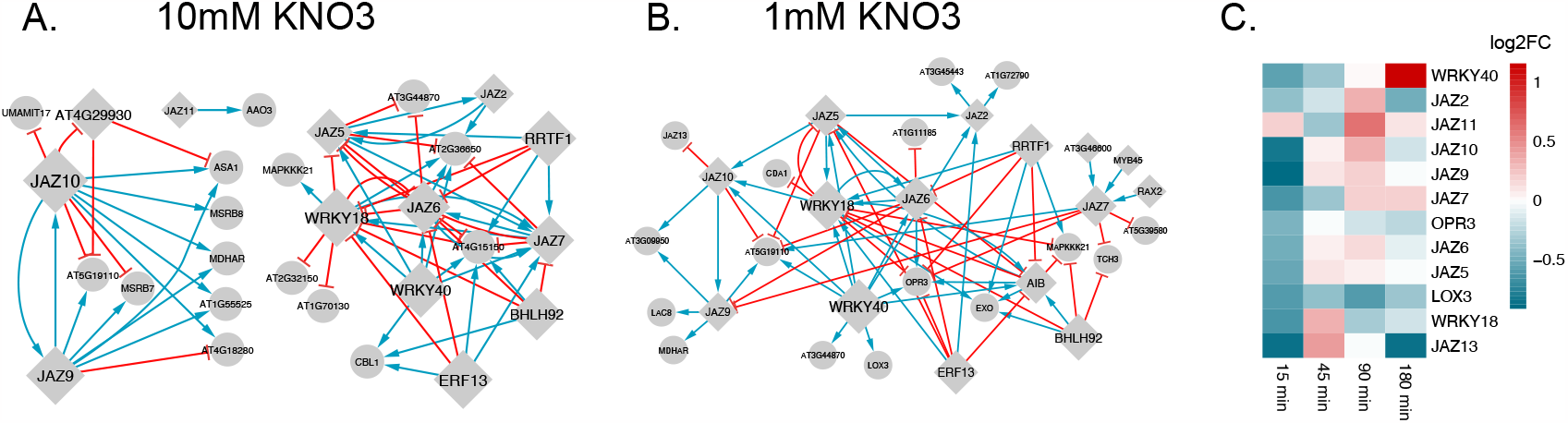
Rewiring of JA-related subnetwork upon KNO_3_ exposure. **A**. 10mM KNO_3_ network. **B**. 1mM KNO_3_ network. **C**. Heatmap of differentially expressed genes linked to JA regulation and metabolism.

### Identification of Key Transcriptional Players in the Nitrogen-Mediated Network Dynamics

Next, we delved into the identification of critical transcription factors (TFs) pivotal for the early transcriptional response to changes in nitrogen (N) availability. Recognizing that TFs play a central role in the temporal rewiring of the network, we embarked on a comprehensive analysis considering network topology, regulatory rewiring, and the impact of TFs on perturbation.^34–36^ To identify these key TFs, we ranked all the TFs taking into account network topology and rewiring. Specifically, we (1) quantified the outdegree of each TF, (2) computed the impact of each TF upon perturbation^37^, (3) and used the DyNet plugin^38^ in Cytoscape to summarize the regulatory rewiring of each TF between the two networks (Table 1). We ranked the TFs by normalizing each metric and, subsequently, taking their sum. Among the most highly ranked genes, we found CCA1, which has previously been shown to link the circadian clock with N metabolism^16^. CCA1 also plays a role in root development: CCA1 overexpression results in altered lateral root architecture.^39^ Within our network, we found that CCA1 participated in 30 interactions in the 1mM network and 20 interactions in the 10mM network, with 13 of these interactions found in both networks. The eighth-ranked gene, SCARECROW-LIKE 28 (SCL28), participated in 13 interactions in the 1mM network and 20 interactions in the 10mM network, with only 1 of these interactions found in both networks. Mutants of *SCL28* showed altered RSA with reduced primary root length.^40^ CCA1 and SCL28 have not been characterized in the context of altered N-dependent responses in terms of root system architecture, nor have nine of the top 10 ranked genes (Supplementary Figure 4).

**Table 1.**
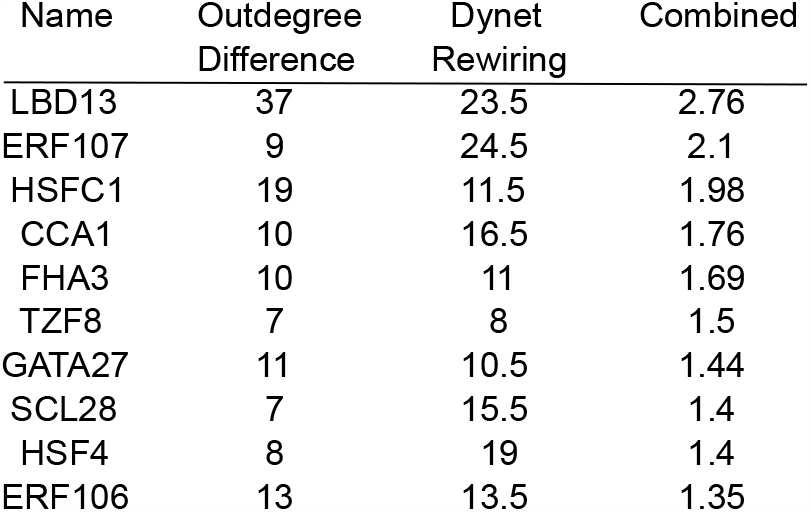

Further exploration into TFs crucial for the early transcriptional response led us to ERF106 and 107. Accordingly, genes that were differentially expressed between limiting and sufficient N conditions at 45 and 90 minutes after transfer formed a module generally clustered around the top ranked CCA1 as well as ERF106 and ERF107. Under nitrate limiting conditions, we inferred extensive overlap in direct downstream targets for ERF106, ERF107 and CCA1 (10 shared targets), ERF107 and CCA1 (5 shared targets), and between ERF106 and CCA1 (5 shared targets) (Figure 4A-B). This redundant regulation is less prominent in the sufficient N network where we found 2 common targets with all 3 TFs, 4 between ERF107 and CCA1, and 6 between ERF106 and CCA1. Overall, the topology of this module, specifically the redundant regulation, may indicate the importance of this module in N limiting conditions. As such, we hypothesized that ERF106, ERF107 and CCA1 played an essential role in N-mediated regulation of RSA. To support this hypothesis, we further investigated ERF107. Inferred downstream targets of ERF107 were important N-related genes such as NRT2.4 and CCA1 in the N limiting network. Indeed, we found that *erf107-1* mutants have reduced lateral root length in both 10mM and 1mM nitrate (Figure 4C),^7^ suggesting that this integrated analysis not only provides insights into the hierarchy of transcriptional responses to N availability but also highlights the pivotal roles of specific TFs.

**Figure 4.**
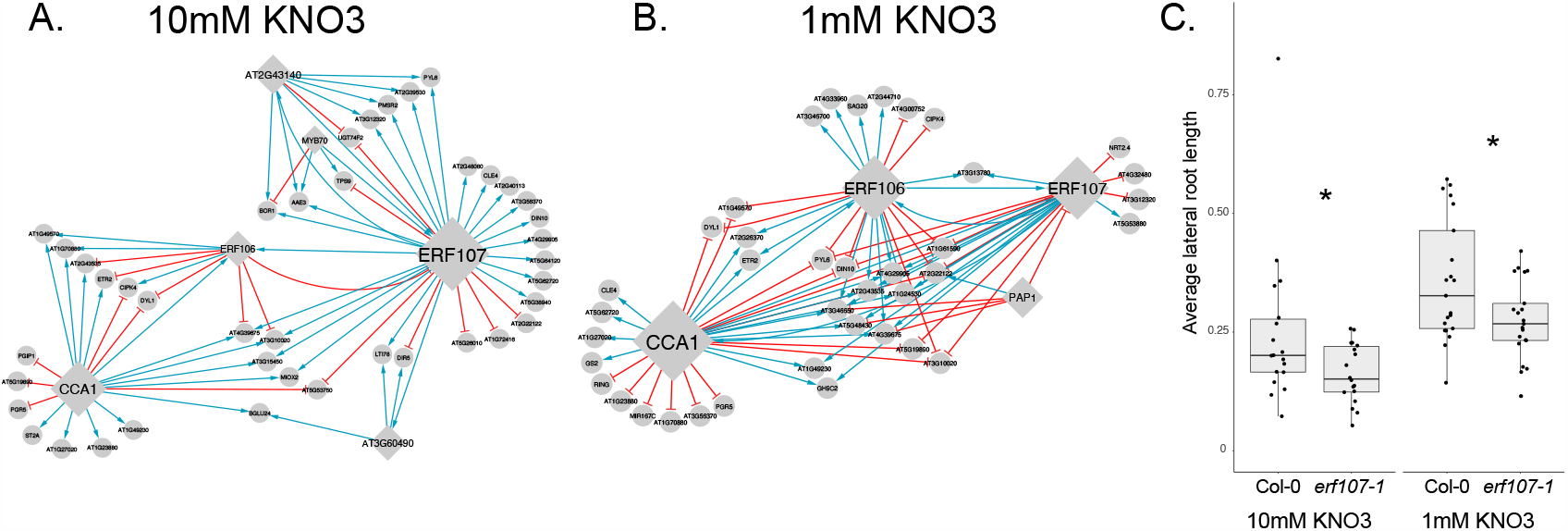
Rewiring of subnetwork linked to ERF107 upon KNO_3_ exposure. **A**. 10mM KNO_3_ network. **B**. 1mM KNO_3_ network. **C**. erf107-1 mutants have significantly shorter average lateral root lengths in 10mM KNO_3_ and 1mM KNO_3_.

### Deciphering LBD13’s Central Role in Nitrogen-Mediated Root Development Dynamics

LBD13 was the top-ranked gene in terms of the number of outgoing targets and its rewiring score (Table 1). Within the inferred network, LBD13 was part of a network module, which consisted of 51 genes (first-degree neighbors) that were primarily responsive to changes in N at the 180 minute time point. Notably, a majority of the regulatory interactions involving LBD13 (40/43) were specific to N limiting conditions (Figure 5B). In N sufficient conditions regulatory interactions for some of these genes are rewired with connections to other top genes: HEAT SHOCK FACTOR C1 (12 genes, 4 genes with the same interaction type), HEAT SHOCK FACTOR 4 (9 genes, 3 genes with the same interaction type), and SCL28 (1 gene with the same interaction type) (Figure 5A-B).

**Figure 5.**
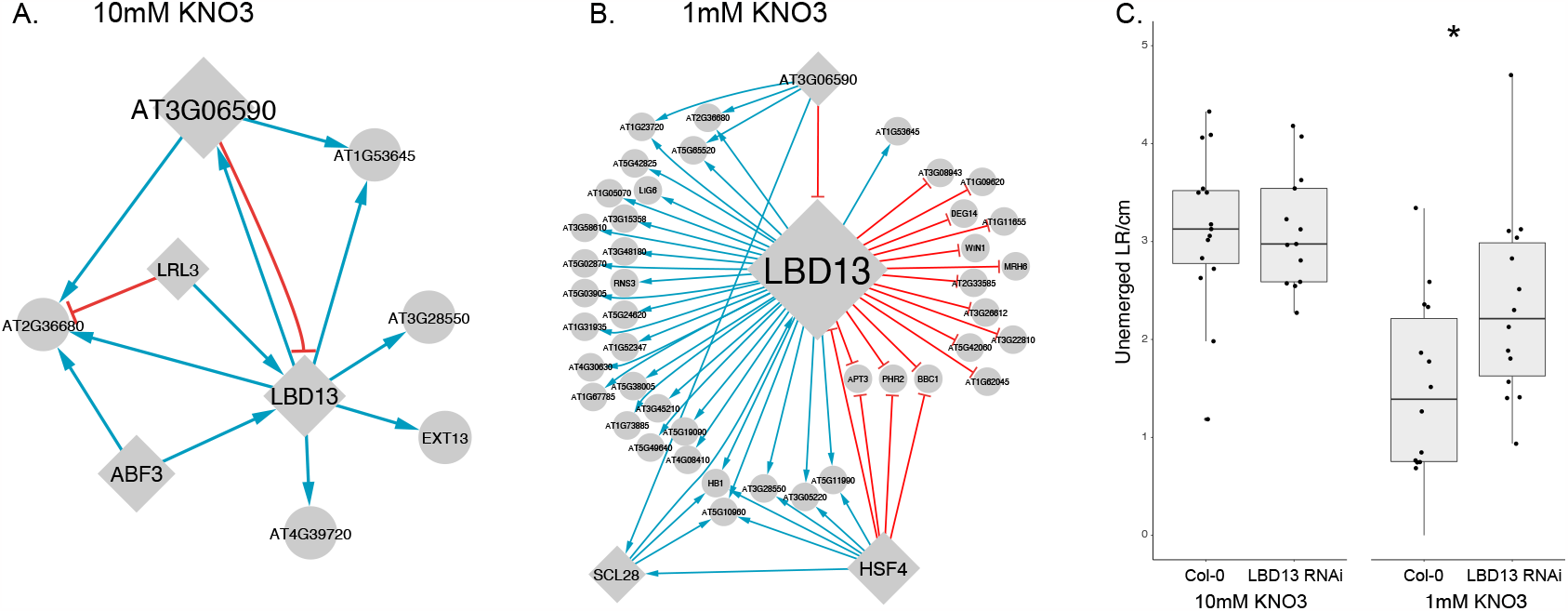
Rewiring of subnetwork linked to LBD13 upon KNO_3_ exposure. **A**. 10mM KNO_3_ network. **B**. 1mM KNO_3_ network. **C**. LBD13 RNAi mutants have significantly more unemerged lateral roots per cm than Col-0 wildtype seedlings.

While LBD13 has previously been shown to regulate lateral root development, its function in response to changes in N availability is unknown. We used this inducible RNAi line of *LBD13*.^41^ and analyzed its RSA in limiting and sufficient N in the presence of dexamethasone. In limiting N, we observed a differential lateral root response upon N limiting and sufficient conditions. The number of unemerged lateral roots per cm primary root were increased in N limiting conditions relative to N-sufficient conditions. (Figure 5C). However, we observed no other RSA-related mutant phenotypes related to primary and lateral roots in the LBD13 RNAi line. In conclusion, in accordance with the network ranking predictions, LBD13 acts as a central regulator by controlling lateral root emergence in a N condition-specific manner. As the LBD13 RNAi line showed differential lateral root emergence in limiting vs sufficient N, we hypothesized that LBD13 may also play a role in early lateral root development. Indeed, *LBD13* had a peak of expression in lateral root founder cells, which declined as the founder cells progressed through asymmetric cell division and auxin response^42^ (Supplemental Table 5). To further dissect the role of LBD13 in early lateral root development, we leveraged cell-type specific single cell expression profiling data that delineates a developmental trajectory associated with lateral root initiation in N-sufficient conditions and used dynamic Bayesian network inference to generate a TF regulatory network (TFRN) from the N-responsive TFs (Supplemental Table 3D, Supplemental Figure 5). Several biologically relevant network motifs were enriched within this early LR initiation network, including bi-fans, feed forward loops (FFLs), feedback loops, and bi-parallels^43^ (Figure 6A, Supplemental Table 6). FFLs have been shown to be at the base of a plant’s adaptation to stimuli^44^ and to play important roles throughout cellular development and growth, such as accelerating an output response and generating pulse-like dynamics.^45^ We therefore further focused on a type I coherent feedforward loop^46^ in which LBD13 participates and for which there is pre-existing evidence of these genes regulating early lateral root development. LBD13 is predicted to regulate PDF2 and ERF107; and PDF2 to regulate ERF107. LBD13 regulates the number of emerged lateral roots per cm primary root (Figure 5C); ERF107 regulates lateral root length (Figure 4C) and PDF2 has been shown to function in lateral root development prior to emergence.^47,48^

**Figure 6.**
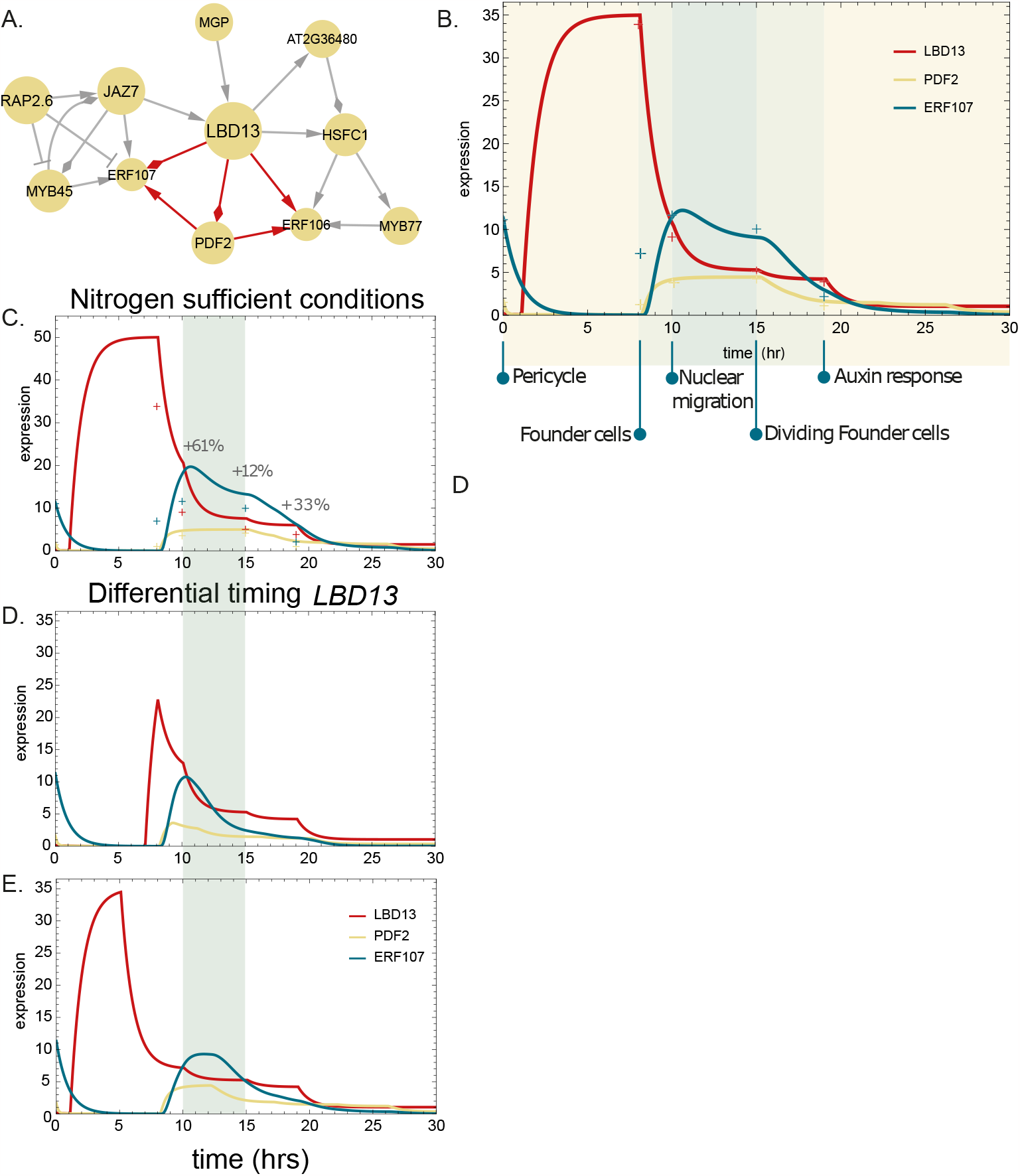
Mathematical modeling of the LBD13 feed forward loop in lateral root development. **A**. The regulatory interactions in which LBD13, PDF2, ERF106, and ERF107 are involved within the N-responsive TF regulatory network during lateral root development. **B**.**-E**. Expression of *LBD13, PDF2*, and *ERF107* modeled in **B**. a wild-type condition, **C**. nitrogen sufficient conditions, **D**.**-E**. and differential timing of *LDB13* during lateral root development. Crosses on the graphs represent the experimental expression values at each development stage.

To analyze the expression dynamics of LBD13 and these two downstream targets in early lateral root initiation, and to link regulatory rewiring to a differential N response, we generated a quantitative model, using ordinary differential equations (ODEs); each of which included a production and degradation term that depended on its respective upstream regulators. We modeled LBD13 expression in a time-dependent manner based on the single cell expression data. In addition, we included the homodimerization of PDF2^48^, which led to a better fit of the model to the expression dynamics of its downstream target ERF107 as observed in the cell-type specific single cell expression data. When modeling a FFL, the two regulators, here LBD13 and PDF2, can act through an OR gate or an AND gate, where only LBD13 or PDF2 would need to be expressed to activate ERF107; or where both TFs need to be expressed to activate ERF107, respectively. The model fits the data better only with an AND gate), suggesting that ERF107’s expression is tightly regulated by multiple factors. Moreover, our model suggests that the specific spatial expression of *LBD13* in the pericycle cells as they transition to founder cells is key for proper lateral root initiation. The model demonstrated that LBD13 in the founder cells induces *PDF2* at the stage of nuclear migration and in dividing founder cells. Together PDF2 and LBD13 activate *ERF107*, potentially inducing nuclear migration and founder cell divisions. To evaluate how the dynamics between LBD13, PDF2, and ERF107 affect lateral root development upon nitrogen limiting conditions, we adjusted *LBD13* expression in our model to its expression observed in our time course data under nitrogen limiting conditions, which is a 43% increase. After running our model under these new conditions, we observed an increased and extended expression domain of PDF2 and ERF107 (Figure 6C).

We explored the robustness of our model by changing the duration of *LBD13* induction. Lower expression of *LBD13* at a later time point resulted in a lower peak of *ERF107* expression and no expression in the dividing founder cells (Figure 5D). In contrast, earlier expression of *LBD13* at the same magnitude as in the original model, but with a shorter duration resulted in a peak of expression in founder cells during nuclear migration (Figure 6E). In each of these cases of modulation of *LBD13* expression magnitude and developmental stage, *PDF2* expression duration was the same (from nuclear migration to the initiation of the auxin response), but its relative magnitude shifted during these time periods (Figure 6C-E). The results show that *LBD13*’s spatiotemporal expression matters for the proper transcriptional progression of PDF2 and ERF107 and thus also potentially for lateral root initiation. In essence, this modeling provides further evidence for the importance of LBD13 in orchestrating a feedforward loop that plays an important role in initiation of ERF107 expression and early events in lateral root initiation. This initiation is generally repressed; only when there are sufficient amounts of *LBD13* is root initiation stimulated and the loop is activated. Upon 10mM nitrogen, *LBD13’*s expression is reduced and does not reach sufficient levels as much, leading to a reduction in lateral roots. Taken together, our study identifies LBD13 as a central regulator in nitrogen-responsive transcriptional networks, finely modulating lateral root dynamics in response to specific nitrogen conditions. This pivotal role advances our understanding of the intricate regulatory mechanisms governing root development, especially in the context of variable nitrogen availability.

## Discussion

In this study, we used a transcriptomic time course of roots exposed to limiting and sufficient nitrate conditions to infer an associated regulatory network. Here, we performed a time course experiment after N addition, profiling expression in roots using growth conditions we previously established to have significantly different effects on RSA.^7^ While several time course experiments profiling aspects of the N response have been published^2,4^, experimental details such as N source and concentration, developmental age, organ type, and time of day of harvest are all relevant factors when interpreting resulting data. All these studies provide complementary datasets that can be mined to discover genes involved in the regulation of N-mediated growth and metabolism.

We analyzed the RNAseq time course data by identifying genes that were differentially expressed between the two N conditions at the same time point. The observed differential expression, therefore, is reflective of the altered N treatment as compared to our starting 0 minute time point. Taking these lists of DEGs and our knowledge of their temporal N responses, we used a machine learning approach with TuxNet to infer regulatory interactions and GRNs for each N condition. We used the network structure and regulatory modules as a guide to identify genes of interest. In comparing the two networks, we found that there were many TFs that have dynamic regulatory roles. These TFs were ranked for their impact on network connectivity and rewiring.

Network-wide enrichment analysis revealed that JA signaling genes are overrepresented in the network. The importance of the JA-related module is illustrated through its prominent size and its considerable rewiring between the N conditions. Although JA has been associated with the regulation of auxin production in roots, and linked through a network mapped through enhanced yeast one hybrid (eY1H) analyses, there have been no direct connections made to link JA responses to N treatment to regulate RSA. These JA-related genes are initially upregulated, likely due to transfer of seedlings to new plates, and are more highly expressed in the limiting N conditions. This indicates that sufficient N in the environment may temper the JA-mediated plant stress response after they were moved to new plates. Collectively, these data demonstrate a transcriptional link between JA mediated signaling and changes in N nutrient status within Arabidopsis roots.

ERF107 was previously identified to bind to the promoters of 22 genes, including prominent N uptake and biosynthesis genes such as NFP6.3/NRT1.1 and NIA1 in eY1H assays. The prominence of ERF107 in both networks and the *erf107* mutant root phenotype support its role as a regulator of N-mediated root responses. Although the 22 ERF107 targets identified in the eY1H-generated network were not identified in this inferred network, it may be due to spatio-temporal or N condition-specific aspects of the response that were not captured in our time course transcriptome. To validate our top ranked TF, LBD13, we performed root phenotyping assays using the same conditions as the time course. The LBD13 RNAi plants displayed altered lateral root development, but only in limiting N conditions. This phenotype supports LBD13’s predicted role in the regulatory networks, where it is central to the structure of a module in the limited N network while having a minimal role in the sufficient N network. To further investigate the role of LBD13 and its predicted targets, ERF107 and PDF2, in lateral root development, we incorporated the dynamic nature of these interactions using ODEs coupled with a single cell transcriptome dataset of very early steps in lateral root initiation. This modeling reinforced our hypothesis that LBD13 acts in the early stages of lateral root development, acting through a feedforward loop with PDF2 and ERF107. This modeling was further used to predict the change in the network when in the presence of limiting relative to sufficient N.

These results demonstrate the power of diverse methods using transcriptome data to map and infer gene regulatory networks to successfully identify genes and regulatory circuits that regulate a process of interest. Using these data as guidance, information from the structure of our limiting and sufficient N networks, in the future we can test the higher order interactions within and between these regulatory modules. Delving into these relationships will reveal emergent properties of the N networks, their structure, and the possible redundancy that is a hallmark of many biological networks.^49,50^

## Materials and Methods

### Plant material and growth conditions

For sequencing libraries, Col-0 seeds were surface sterilized and plated on mesh on 1mM KNO_3_ media (recipe found in Gaudinier et al 2018^7^) and stratified for 2 days at 4°C. Plates were then placed vertically and grown in 22°C in 16 hour days/8 hour nights. 7 day old plants were transferred either to new 1mM or 10mM KNO3 plants at two hours post-dawn in the growth chamber. Per biological replicate, two plates per time point were transferred then combined. Root tissue was then harvested and flash frozen in liquid N_2_ at 0 min (mesh was picked up and placed down on the same plate to account for mechanical responses to transfer), 15 min, 45 min, 90 min, and 180 min. We performed four biological replicates for the time courses.

For RSA phenotyping, the LBD13 RNAi seeds were generously shared with us by Jungmook Kim^41^. Col-0 and LBD13 RNAi seeds were surface sterilized and plated on 1mM KNO_3_ or 10mM KNO_3_ media containing 10 μM dexamethasone and were stratified for 2 days at 4°C. Plates were then placed vertically and grown in 22°C in 16 hour days/8 hour nights. Plants were grown for 7 days and imaged using a light box and a Canon EOS Rebel T7. Lateral root stages (emerged and unemerged) were quantified using a Nikon Diaphot TMD inverted microscope. Primary roots of 9-day-old seedlings were traced using a Wacom Bamboo tablet in ImageJ. Data were log-transformed and analyzed using ANOVA in R.

### RNA-seq library preparation and pooling of technical replicates

RNA-seq libraries were prepared following the BRAD-Seq DGE protocol.^51^ Libraries were sequenced using the Illumina HiSeq 3000 in SR50 mode.

### Transcriptome analysis and Network inference

Reads of each sample were mapped against the *Arabidopsis thaliana* reference genome (TAIR v. 10) with TuxNet^22^ using default settings. TuxNet specifically uses ea-utils fastq-mcf for preprocessing^52^, hisat2 to align the reads to the reference genome^53^, and Cufflinks for differential expression analysis^54^. To identify differentially expressed genes (DEGs), pairwise comparisons between both treatments at each time point were performed using an FDR < 0.05. Heatmaps and plots of the DEGs were generated in R (version 4.0.2) using ggplot2^55^.

To infer gene regulatory networks, we first selected the short-term DEGs (i.e. DEGs at 15m, 45m, 90m, and 180m) as input set. Next, using the FPKM replicate values of the limited nitrogen dataset, we inferred a regulatory network between our input DEG set with a random forest approach (RTP-STAR) within the TuxNet interface. The regulatory interactions between the same set of DEGs were inferred within the sufficient nitrogen network by using the FPKM replicate values from the 10mM KNO_3_ time course in the machine learning approach. Only putative TF-encoding genes were considered as source nodes that could regulate the expression of other DEGs. To infer a nitrogen-responsive network during lateral root development, the 82 nitrogen-responsive TFs were selected. Using Cabrera Chavez et al^42^ cell-type specific expression dataset, we inferred a TF regulating network with Bayesian principles. Specifically, we used GENIST from the TuxNet interface. As settings we used Reg Time Percent 0.5, Reg Fold Change Threshold 1.25, and Time Lapse 0 and 1. TuxNet is available at https://github.com/rspurney/TuxNet. Network visualizations were made using Cytoscape v3.8.0.^56^

### Network Analyses

#### Gene Ontology Enrichment

Enriched GO terms were identified with PANTHER. To summarize and reduce redundant GO terms, Revigo^57^ was used. The GO enrichment plot was generated in R (version 4.2.2) using ggplot2^55^.

#### DAP-seq

Binding analysis data was downloaded from http://neomorph.salk.edu/dap_web/pages/index.php and all regulators and their targets were searched for using a custom R script.

#### N and JA enrichment

To test for enrichment of genes in the network using N-related datasets, Fisher’s exact test was used in R, using the standard function fisher.test(). For this, we used the gene list from Table 2 from Canales *et al* 2014^20^, Supplementary Table 2 from Liu *et al* 2017^21^, and Supplementary Table 3a from Gaudinier *et al* 2018^6^. We tested whether the overlap between differentially expressed genes within the inferred network was greater than differentially expressed genes which did not overlap with genes in the network.

To test for enrichment of genes in the network for jasmonic acid related genes, we performed the same fisher.test() using the JA-responsive genes from Zhang et al^23^ Table S2.

### Mathematical modeling

The mathematical model consisted of two ordinary differential equations (ODEs) for PDF2 and ERF107, and a time-dependent equation for LBD13. The interaction of PDF2, ERF107, and LBD13 were formulated as a type 1 coherent feed-forward loop with an AND gate. The spatial expression across lateral root development of LBD13 was modeled time-dependent. The expression of PDF2 and ERF107 is under the control of LBD13, and LBD13 and PDF2, respectively. The regulatory interactions between these proteins were modeled using Hill equation dynamics. For the ODEs, it was assumed that transcription and translation happen quickly, such that transcription and protein degradation could be modeled in the same equation. All proteins are assumed to have a linear degradation term.

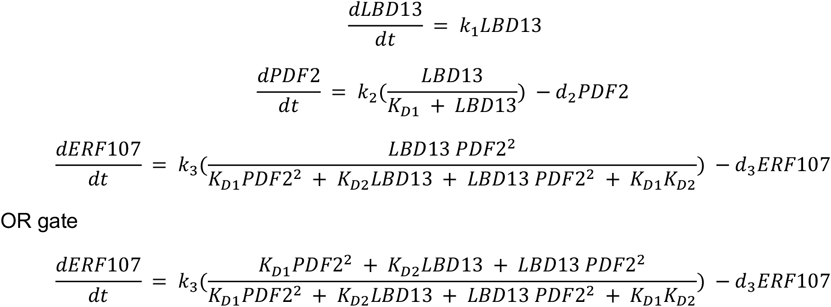

Model simulation was done with Mathematica (Wolfram, Inc., Champaign, IN). The source code for the equations, model simulations, and plotting are provided on https://github.com/LisaVdB/LBD13.

## Acknowledgements

We thank J. Kim for LBD13 RNAi seeds. We thank M. Tang, G.M. Turco, and A.-M. Bagman for help with the time course, B.K. Blackman for laboratory and growth chamber space for root growth assays and D.J. Kliebenstein for manuscript discussions. We gratefully acknowledge the support of our funding sources. This work was funded by DuPont Pioneer and the Miller Institute for Basic Research in Science Postdoctoral Fellowship awarded to A.G. This work was funded by a joint USDA/NSF/BBSRC Breakthrough Technologies Award Grant numbers: USDA: 2019-67013-29012, BBSRC: BB/S020853/1) to S.M.B. Additional funding was provided by NSF 2118017 and 2119820 and a Howard Hughes Faculty Scholar award to S.M.B. This work was supported by the Foundation for Food and Agriculture Research (FFAR CA18-SS-0000000026), Benson Hill, VIB, BASF, the United Soybean Board (2020-152-0134), and the North Carolina Soybean Producers Association (20-122) and the National Science Foundation (NSF) (PGRP BIO-2112058) to R.S. This work was funded by Ministerio de Ciencia e Innovacion (MICIN) of Spain and ERDF (grants PID2019-111523GB-I00 Facturas_PID2022-140719NB-I00 to M.A.M.-R.) and by the ‘Severo Ochoa Program for Centres of Excellence in R&D’ from MCIN/AEI /10.13039/501100011033 (grant CEX2020-000999-S) to CBGP.

